# Investigating lead removal at trace concentrations from water by inactive yeast cells

**DOI:** 10.1101/2021.10.07.463380

**Authors:** Patritsia M. Stathatou, Christos E. Athanasiou, Marios Tsezos, John W. Goss, Camron Blackburn, Filippos Tourlomousis, Andreas Mershin, Brian W. Sheldon, Nitin P. Padture, Eric M. Darling, Huajian Gao, Neil Gershenfeld

**Author notes:** These authors contributed equally to this work (Patritsia M. Stathatou, Christos E. Athanasiou).

## Abstract

Traces of heavy metals found in water resources, due to mining activities and e-waste discharge, pose a global threat. Conventional treatment processes fail to remove toxic heavy metals, such as lead, from drinking water in a resource-efficient manner when their initial concentrations are low. Here, we show that by using the yeast *Saccharomyces cerevisiae* we can effectively remove trace lead from water *via* a rapid mass transfer process, achieving an uptake of up to 12 mg lead per gram of biomass in solutions with initial lead concentrations below 1 part per million. We found that the yeast cell wall plays a crucial role in this process, with its mannoproteins and β-glucans being the key potential lead adsorbents. Furthermore, we discovered that biosorption is linked to a significant increase in cell wall stiffness. These findings open new opportunities for using environmentally friendly and abundant biomaterials for advanced water treatment targeting emerging contaminants.

**One-Sentence Summary:** Removing toxic heavy metals from water at challenging trace levels in an environmentally friendly, resource-efficient manner.

---

Heavy metals are highly water-soluble and non-biodegradable, tending to persist indefinitely when released into water bodies. Electronic waste (e-waste) discharge and mining are the most dominant anthropogenic activities responsible for heavy metal contamination of water resources(*1*). Acid mine drainage (AMD), *i*.*e*., leakage of highly acidic water rich in metals, is a global environmental threat(*2*). In the United States (US) alone, AMD is the main source of water pollution, impacting currently over 20,000 km of streams(*3*), deriving from the 13,000 active and the 500,000 abandoned mines(*4*), which continue generating AMD for centuries after their closure(*5*). In addition, around 50 million tons of e-waste was globally produced in 2018, expected to reach 120 million tons per year by 2050. Over 80% of this e-waste either ends up in landfills or is being recycled under poorly regulated conditions, using primitive and pollutive methods, hence, seriously contaminating water resources(*6*).

Lead (Pb) is one of the most widely used heavy metals; its production increased by about 20% during the last decade, reaching around 11.7 million tons globally in 2020(*7*). Pb is highly toxic, even at trace concentrations, with deleterious effects on organs and tissues of the human body(*8*). It can enter drinking water either due to inadequate water treatment, or due to chemical reactions with Pb-containing components of water distribution systems(*9, 10*). After numerous incidents of Pb contamination(*9, 11*), with most recent the water crisis in the city of Flint, Michigan, USA in 2014, limits of Pb in drinking water are becoming more stringent: in 2018, the European Commission proposed reducing Pb limits to 5 parts per billion (ppb)(*12*), while in 2020, the US Environmental Protection Agency determined that no level of Pb in drinking water is safe(*13*).

Conventional water treatment methods either fail to completely remove trace Pb amounts or result in significant financial and environmental costs to do so(*14*–*16*). Biosorption, a mass transfer process by which heavy metal ions bind onto inactive biological materials by physicochemical interactions, is a competitive alternative to conventional processes, as abundant biomass sources can be effective, practical, and sustainable adsorbents(*17*). Although biosorption of heavy metals has been studied at the parts per million (ppm) contaminants scale, there is a significant research gap at the challenging trace initial concentrations of ppb and below.

In this study, the unexplored Pb biosorption mechanisms at the ppb scale are investigated using inactive yeast biomass as the biosorbent. A strain of the common yeast, *Saccharomyces cerevisiae (S. cerevisiae)*, was selected, as it is a biodegradable adsorbent, widely used in various industrial settings(*15, 18*), that effectively remove Pb at ppm initial concentrations(*19*). The yeast cells were harvested at the peak of their exponential growth phase (Fig. 1A), for optimal biosorptive capacity(*19*). The harvested cells were washed to remove culture medium residues and metabolites (Fig. 1B), before being lyophilized (freeze-dried) and converted to powder (Fig. 1C). Kinetic and equilibrium experiments were conducted by adding this yeast powder biomaterial to ultrapure water spiked with lead(II) nitrate (Pb(NO_3_)_2_). After the required contact time, liquid and solid phases were separated and analyzed to measure residual Pb concentrations and identify potential biomass sites responsible for Pb uptake (Fig. 1D). Overall, this work showcases the use of an effective trace heavy-metal removal biomaterial, made from an environmentally friendly, inexpensive, benign to human health, and easy-to-mass-produce microorganism.

**Fig. 1.**
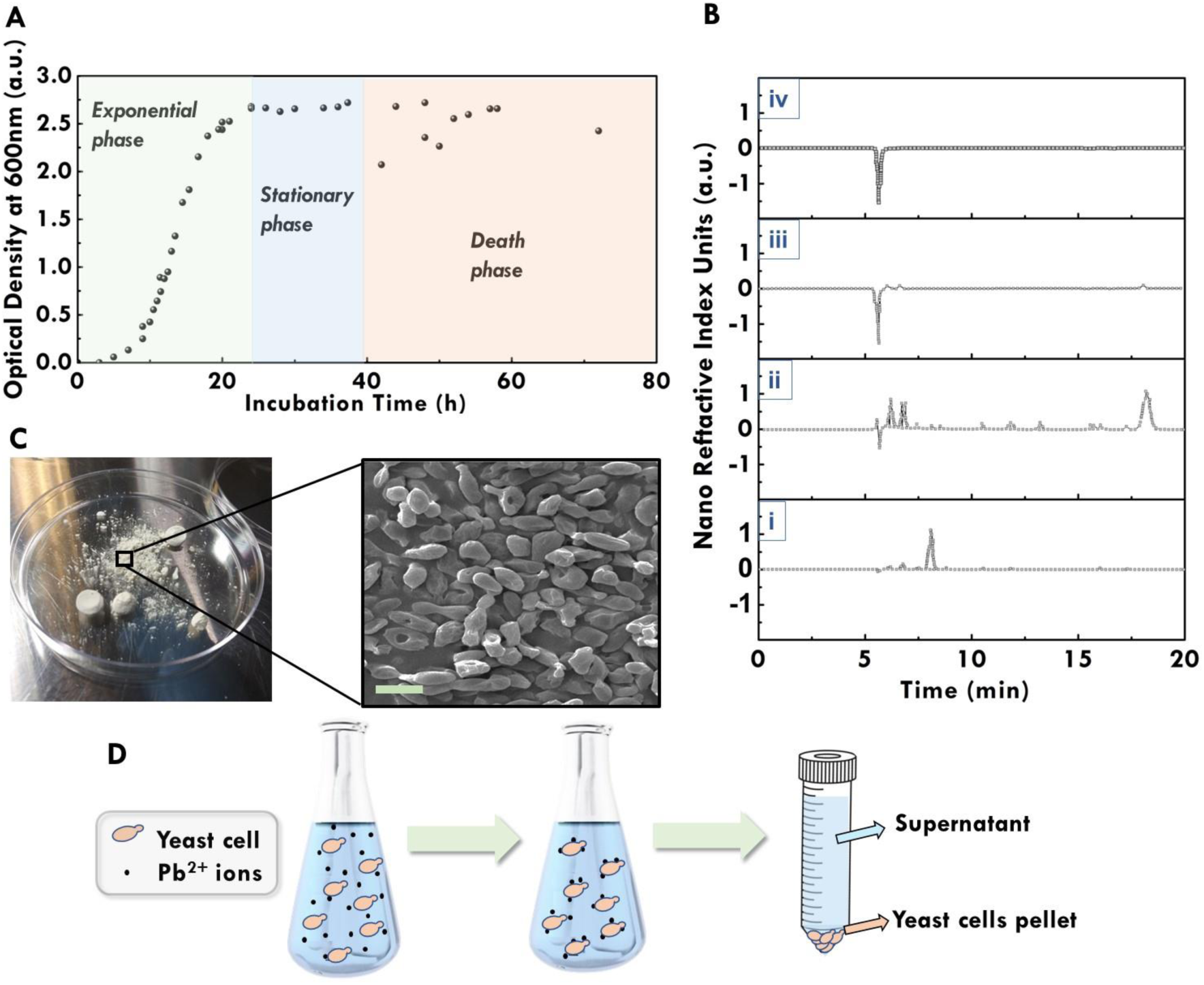
Overview of experimental procedures. (**A**) Measured growth curve of *S. cerevisiae* yeast cells (absorbance accuracy is approximately 1.75% of reported values). (B) High-performance liquid chromatography (HPLC) analysis results, performed to identify the number of washes with ultrapure water required to remove medium residues and metabolites from harvested yeast cells before being used in biosorption experiments: i) pure culture medium showed a peak at 8 mins representing glucose; ii) in the supernatant of the harvested liquid culture, glucose was not present but ethanol was produced (peak at 18 mins); iii) in the supernatant after the first wash of yeast cells with ultrapure water there were almost no peaks representing the presence of sugars or organic acids in the solution; iv) in the supernatant after the second wash of yeast cells with ultrapure water there were no peaks at all. Therefore, it was decided to wash the harvested cells twice to ensure absence of organic compounds that might affect further experiments. (C) Yeast powder after lyophilization and scanning electron microscope (SEM) imaging of freeze-dried yeast cells (scale bar: 5 μm). (D) Main steps of kinetic and equilibrium experiments involving the addition of freeze-dried yeast cells in Pb-containing aqueous solutions, the adsorption of Pb ions, and the separation of biomass and supernatant after the required contact time *via* centrifugation for further analyses.

## Results

### Effect of solution pH on Pb speciation and uptake

Solution pH is a key parameter for biosorption, as it affects the chemistry and speciation of both the metal-uptaking functional groups in the biomass, and of the hydrolyzed Pb ionic forms. Pb speciation in the solution is also affected by the Pb concentration at any given pH and oxidation state(*17, 20*). To quantify the resulting Pb speciation after the hydrolysis of Pb(NO_3_)_2_ in a wide pH range (3-13), at 25 °C, and at a given initial concentration (*C*_*0*_) of 1 μM Pb(NO_3_)_2_, the Eawag ChemEQL v3.2 software(*21*) was used. Pb^2+^ is the dominant species until pH reaches 5.8, where lead hydroxides (*e*.*g*., PbOH^+^) are beginning to form (Fig. 2A).

**Fig. 2:**
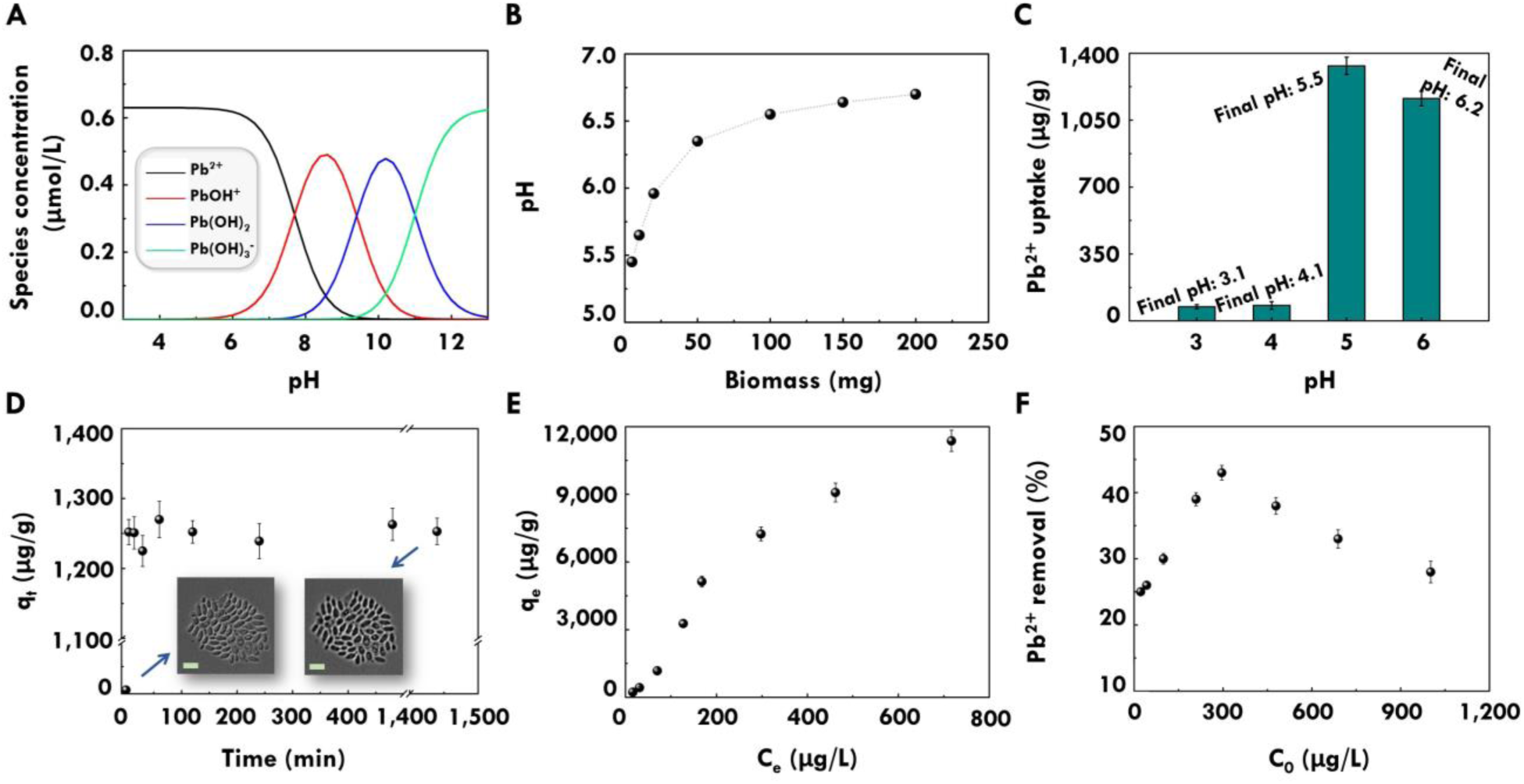
Effect of solution pH on Pb speciation & uptake, biosorption kinetics, and adsorption isotherm. (**A**) Distribution of Pb(NO_3_)_2_ hydrolysis products at 25°C and 1 μM Pb(NO_3_)_2_. (B) Increase in solution pH due to yeast biomass addition; error: ±0.06 pH units. (C) Effect of initial pH of solution on Pb^2+^ uptake for 5 mg of yeast biomass and *C*_*0*_ 100 ppb Pb^2+^. The final pH of solutions after biosorption is reported for each case; error: ±0.06 pH units. (D) Kinetic experiment of 5 mg yeast biomass with *C*_*0*_ 100 ppb Pb^2+^, indicating the rapid biosorption process; *q*_*t*_: Pb^2+^ uptake capacity of yeast biomass at different time intervals (μg of Pb^2+^ by g of biomass). Time course micrographs of lyophilized *S. cerevisiae* cells incubated in the same aqueous solution at 0 h and 24 h are shown in the inset, validating that there is no cell growth or division during the experiments (scale bar: 8 μm). (E) Adsorption isotherm at 25°C, following the Langmuir adsorption isotherm model; *q*_*e*_: Pb^2+^ uptake capacity of yeast biomass at equilibrium (μg of Pb^2+^ by g of biomass). (F) Pb^2+^ percentage removal versus Pb^2+^ *C*_*0*_.

Solution pH increases with the addition of biomass and plateaus for biomass values greater than 100 mg (Fig. 2B). Therefore, to assess the effect of solution pH on the biomass Pb^2+^ uptake capacity (*q*, in μg of Pb^2+^ by g of biomass), the initial pH of the aqueous solution (before biomass addition) was adjusted to pH values within the range of 3-7, and then 5 mg of yeast biomass were added to moderate the anticipated pH increase due to biomass addition. Pb^2+^ concentrations and pH values were measured both before biomass addition and after Pb^2+^ biosorption (contact time of 24 h). The biomass Pb^2+^ uptake capacity increased significantly as the initial solution pH was increased from 3 to 5 (Fig. 2C). For pH ≥ 6.0 the measured initial Pb^2+^ concentration was significantly lower than the known amount added to the solutions, indicating loss of soluble Pb analytes due to precipitation. This is validated by the formation of lead hydroxides after pH 5.8 (Fig. 2A). The increase of solution pH by biomass addition, as well as the increase of *q* with increasing pH values could be attributed to a potential protonation of the functional groups of yeast biomass at pH values below the pH point zero, *i*.*e*., the pH at which the overall biomass surface charge is zero. At pH values below the pH point zero, the biomaterial may exhibit an overall positive charge, thus attracting negatively charged species and not adsorbing Pb^2+^cations resulting in lower *q* values, while at higher pH values the biomass surface could acquire negative charges leading to increased Pb^2+^ uptake. However, such an approach assumes a rather simplistic electrostatic attraction driving mechanism, which has been shown not to be the only case in biosorption(*17*).

Based on these results, pH 5 was proven to be the most suitable for Pb biosorption, where soluble Pb^2+^ is the most dominant species in the solution and *q* is maximized. All experiments described in the following sections were performed by adding 5 mg of yeast biomass (unless otherwise specified) in aqueous solutions, after adjusting their initial pH to 5 at 25 °C, agitated at 200 rpm.

### Adsorption kinetics & growth analysis of lyophilized yeast

Kinetic experiments were conducted to determine the change in Pb^2+^ concentration in the liquid phase as a function of contact time and identify the contact time required to attain equilibrium. Lyophilized yeast cells were added in aqueous solutions with *C*_*0*_ of 100 ppb Pb^2+^ and were incubated for 24 hours. Samples were taken at specific time intervals (*i*.*e*., 0 min, 5 min, 15 min, 30 min, 1 h, 2 h, 4 h, 8 h, 24 h) and analyzed using inductively coupled plasma mass spectrometry (ICP-MS). As observed, biosorption is a rapid process, with equilibrium being reached within the first five minutes of contact (Fig. 2D). Kinetics data fit accurately with the pseudo-first-order model(*22*), (R^2^: 0.99), with the pseudo-first-order rate constant (*k*_*1*_) equal to 111.98 h^-1^.

In parallel the freeze-dried *S. cerevisiae* cells were incubated for 24 h under conditions identical to the kinetic experiments and observed by phase-contrast microscopy, while acquiring optical density (OD) measurements, to validate that the cells remain inactive during biosorption. Indeed, after 24 h there was neither cell growth nor cell division observed (Fig. 2D); time-lapse video and further information confirming these results are included in the Supplementary Materials of this work.

### Adsorption isotherm

Aqueous solutions with different initial Pb^2+^ concentrations (*C*_*0*_: 20, 40, 100, 200, 300, 500, 700, and 1,000 ppb) were incubated for 1 h. The equilibrium Pb^2+^ concentrations (*Ceq*) were measured after biosorption, and the Pb^2+^ uptake capacity of yeast biomass at equilibrium (*q*_*e*_) was calculated (using Equitation 1 of Supplementary Materials) to develop the adsorption isotherm (Fig. 2E). The maximum *q*_*e*_ measured is equal to 12 mg/g, for aqueous solutions with *C*_*0*_of 1,000 ppb Pb^2+^. The experimental data are accurately fitting with the Langmuir adsorption isotherm model(*23*) (R^2^: 0.98), with the ratio of the adsorption and desorption rates (*K*_*L*_) equal to 1.5 L/mg, and the maximum estimated adsorption capacity (*q*_*m*_) equal to 21 mg/g. However, this fit cannot provide any meaningful insight into the biosorption mechanism(*17, 24*).

Pb^2+^ percentage removal versus Pb^2+^ *C*_*0*_ was also measured (Fig. 2F). Pb^2+^ removal for *C*_*0*_ 20 ppb Pb^2+^ is approximately 25% and increases with the increase of *C*_*0*_, reaching a maximum of 43% at *C*_*0*_ 300 ppb Pb^2+^. After this point, Pb^2+^ removal decreases gradually with the decrease of *C*_*0*_, indicating that the optimum uptake capacity of the yeast biomass quantity (5 mg) is reached around *C*_*0*_ 300 ppb Pb^2+^.

### Yeast biomass imaging

Extracellular and intracellular imaging of yeast cells was performed to observe potential changes in their structure after biosorption, using SEM and transmission electron microscope (TEM) imaging, respectively. Yeast cells harvested from ultrapure water (*C*_*0*_: 0 ppb Pb^2+^), served as control cells and were compared with yeast cells harvested from aqueous solutions of *C*_*0*_ 100 ppb Pb^2+^. No morphological change was observed in the yeast cells after Pb^2+^ biosorption, with the structure and dimensions of the cell wall and cytoplasm remaining the same (Fig. 3A, B). Yeast cell walls were ∼180 nm thick, which is the typical cell-wall thickness of the yeast strain used(*25*). However, yeast cell walls became more electron-dense after Pb^2+^ biosorption, indicating the binding of Pb^2+^ ions on them, and in particular on the outer part of the cell wall (Fig. 3C, D).

**Fig. 3:**
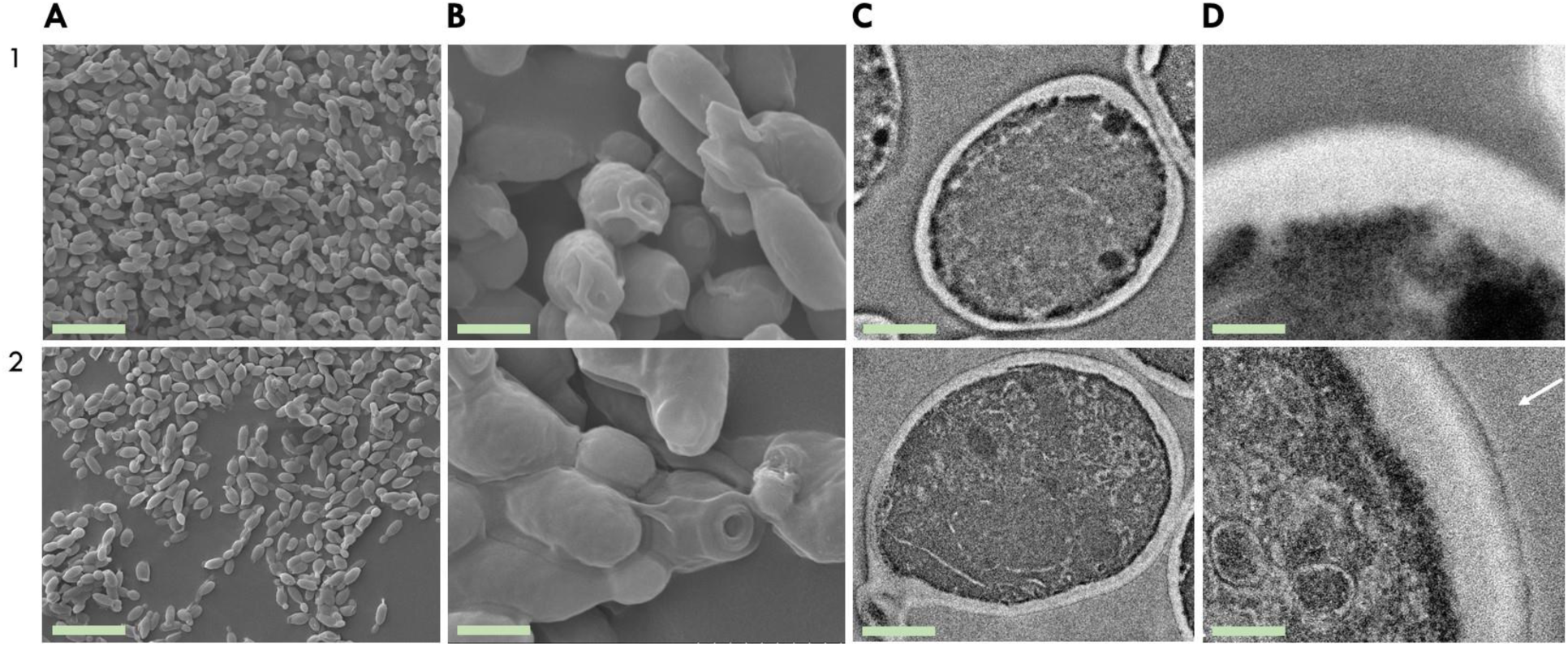
SEM and TEM imaging of yeast cells. Rows are ordered as follows: control yeast cells (*C*_*0*_: 0 ppb) (1), yeast cells after Pb^2+^ biosorption (*C*_*0*_: 100 ppb) (2). (**A**) SEM images presenting an overview of yeast cells (scale bar: 10 μm). (B) Magnified SEM images showing individual yeast cells (scale bar: 2 μm). (C) TEM images of individual yeast cells (scale bar: 1 μm). (D) Magnified TEM images of yeast cell walls (scale bar: 200 nm). The white arrow refers to the outer part of the cell wall, highlighting electron-dense Pb^2+^-binding areas.

### Yeast biomass spectroscopy

Metal biosorption is thought to occur through interactions with functional groups native to the biomass cell wall(*26*). Attenuated-total-reflectance enhanced Fourier transformed infrared spectroscopy (ATR-FTIR) was performed to identify functional groups present in the yeast cell wall and detect changes in them after biosorption, indicating their involvement in Pb^2+^ adsorption. Freeze-dried control (*C*_*0*_: 0 ppb Pb^2+^) and Pb^2+^-exposed yeast cells (*C*_*0*_: 100 and 1,000 ppb Pb^2+^) were analyzed using ATR-FTIR (Fig. 4A). Changes were observed after biosorption in peaks representing C≡N and C≡C stretches, while peak shifts were detected corresponding to N-H in-plane bending from secondary protein amides, which overlaps with the C–N and NO_2_ asymmetric stretching, to vibrational changes of the C-N amide group, and to C–O stretching in the esters and carboxylic acid groups. These changes indicate the contribution of amide and carboxylic acid groups to Pb^2+^ biosorption and the potential role of N in the yeast cell wall on Pb^2+^ binding.

**Fig. 4:**
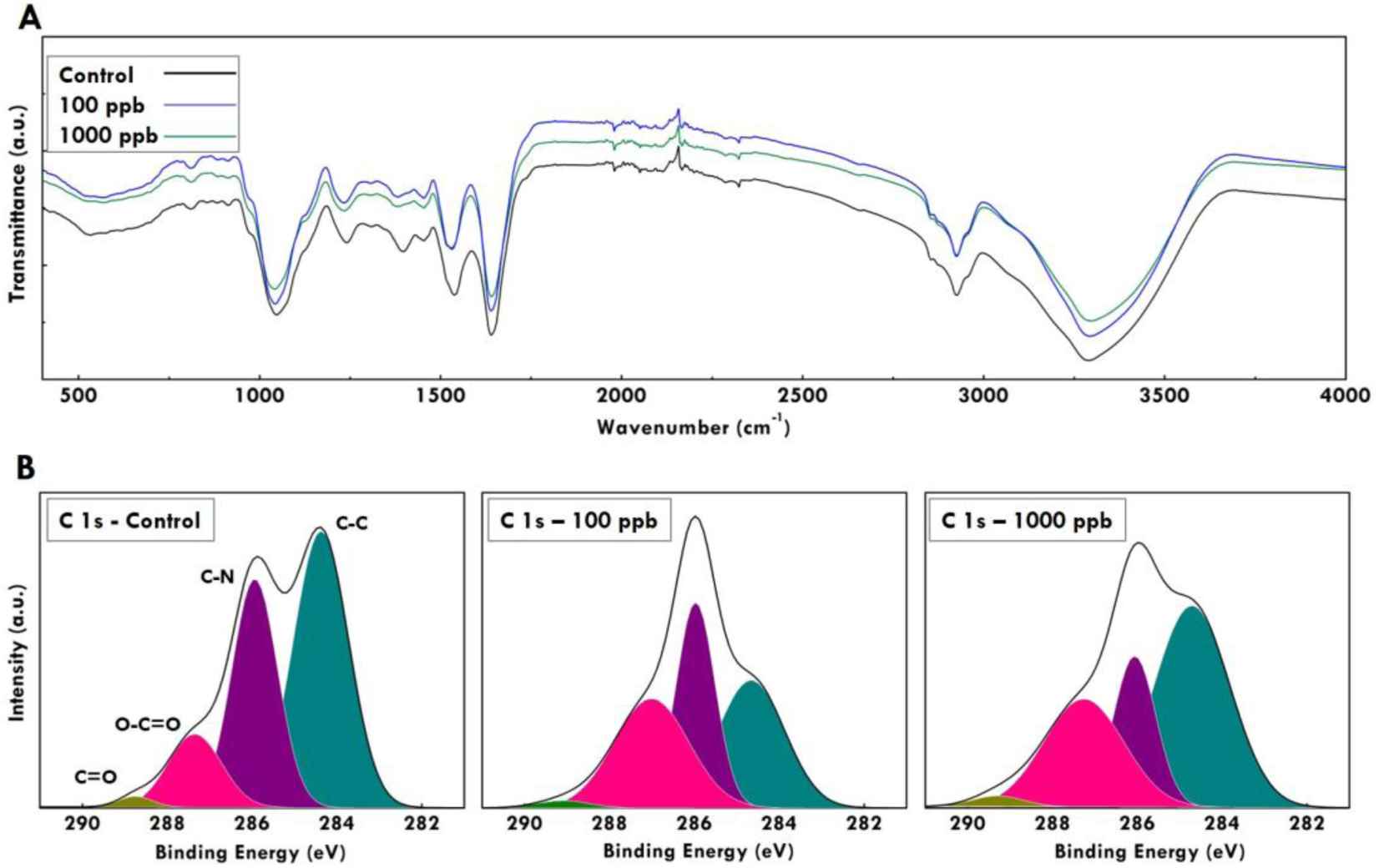
ATR-FTIR and XPS characterization of yeast biomass. (**A**) Full ATR-FTIR spectrum of yeast cells. The black line represents control yeast cells (*C*_*0*_:0 ppb Pb^2+^); the blue and green lines represent yeast cells after biosorption with *C*_*0*_100 and 1,000 ppb Pb^2+^ respectively. (B) XPS analysis of yeast cells. From left to right: control cells, yeast cells after biosorption, *C*_*0*_ 100 and 1,000 ppb Pb^2+^. XPS peak assignments are as follows: light green C=O; magenta: O-C=O; purple: C-N; dark green: C-C.

In addition, the chemical composition of the yeast surface before (*C*_*0*_: 0 ppb Pb^2+^) and after biosorption (*C*_*0*_: 100 and 1,000 ppb Pb^2+^) was analyzed by X-ray photoelectron spectroscopy (XPS) to further explore potential changes in the functional groups of the yeast cell walls. The yeast surface is mainly composed of carbon (C), oxygen (O), and nitrogen (N)(*27, 28*).

Therefore, C 1*s*, O 1*s*, N 1*s* core levels spectra were recorded together with the Pb 4*f*. Significant changes among the control and the Pb^2+^-exposed yeast were only observed for the C 1*s* spectrum, while changes in the Pb spectra could not be detected as the analyzed trace Pb^2+^ concentrations were below the detection limit of the instrument. Deconvolution of the C 1*s* spectrum into Gaussian-shaped lines was performed to identify possible chemical bonds between C, O, and N (Fig. 4B). In all three samples, the C 1*s* peaks are decomposed to peaks at 284.3 eV, 285.9 eV, 287.5 eV, and 288.6 eV representing C-C (*sp*^3^ C), C-N, O-C=O, and C=O respectively. The magnitude and shape of all observed bonds have radically changed after Pb^2+^ biosorption, particularly for C-C, C-N, and O-C=O bonds. These changes indicate the contribution of carboxylic acid and amide groups on Pb^2+^ adsorption, which is consistent with the ATR-FTIR results.

### Chitin’s contribution to Pb^2+^ adsorption

The above analyses indicate that the cell wall of *S. cerevisiae* plays a vital role in Pb^2+^ biosorption. The yeast cell wall has a complex macromolecular structure with a layered organization, including an amorphous inner and a fibrillar outer layer(*29*). The inner layer mainly consists of β-glucans and chitin. The outer layer consists predominantly of mannan polymers, highly glycosylated and linked to proteins (mannoproteins)(*30*).

Several sources suggest that the chitin amine nitrogen is responsible for heavy metals sequestering at the ppm scale(*17, 31, 32*). To further investigate this, we assessed the Pb^2+^ uptake capacity of chitin from shrimp shells, which is similar to that found in the yeast cell wall. We used 20 times more chitin, *i*.*e*., 1.8 mg, than the maximum equivalent amount present in the 5 mg of yeast, considering a 30% dry weight of yeast cell wall and a 6% contribution of chitin per mass to it(*30*). We added this amount to an aqueous solution of 0.2 L with *C*_*0*_ 1,000 ppb Pb^2+^ for 24 h at 200 rpm and 25°C. We ran the same experiments with 5 mg of yeast biomass and with 5 mg of chitin. It was shown that chitin’s Pb^2+^ uptake is negligible, as *C*_*0*_ was reduced by less than 0.3% when we added the 1.8 mg of chitin, which is within the measurement error. Even when the chitin amount added was equal to the total yeast mass (5 mg), *C*_*0*_ was only reduced by 3%, compared to the ∼30% reduction achieved by the yeast biomass (Fig. 5A). Hence, it can be concluded that chitin alone is not contributing to the Pb^2+^ biosorption process.

**Fig. 5:**
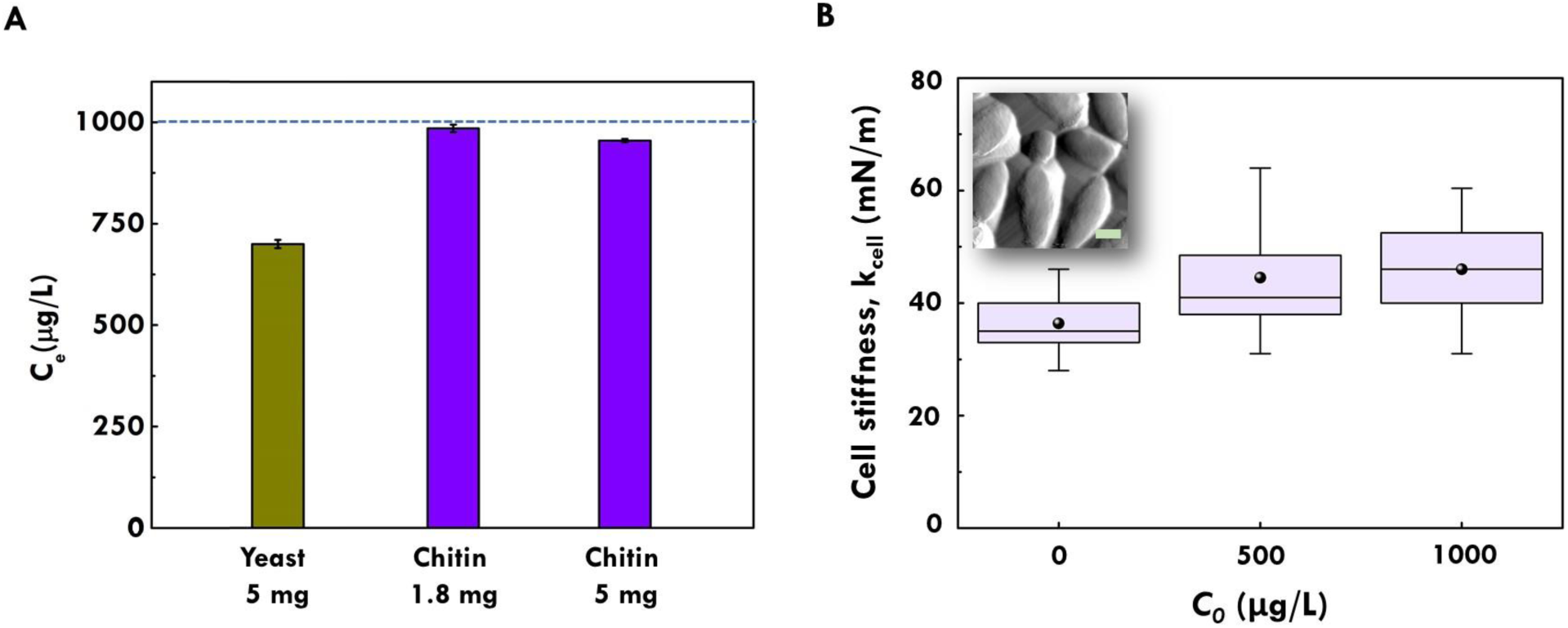
Chitin’s contribution to Pb^2+^ adsorption and nanomechanical characterization of yeast biomass. (**A**) Pb^2+^ *C*_*e*_ after biosorption experiments with 5 mg of yeast biomass (green bar), 1.8 mg, and 5 mg of chitin (purple bars); blue dotted line represents *C*_*0*_: 1,000 ppb Pb^2+^. (B) AFM analysis of yeast cell stiffness. Box-and-whisker plots of the cellular spring constant (k_cell_) calculated from force-extension curves acquired from yeast cells incubated with *C*_*0*_ of 0, 500, and 1,000 ppb Pb^2+^. Boxed region indicates upper and lower quartiles for each data set; median is indicated by the horizontal line within the box; mean is indicated by the bullet point; whiskers extend to high and low data points (n > 70 force measurements from ≥ 8 cells per condition). AFM deflection retrace image of yeast cells is shown in the inset (scale bar: 2 μm).

### Yeast biomass nanomechanical characterization

We employed nanomechanical characterization to investigate biosorption. We assessed the stiffness of the yeast cells before and after Pb^2+^ exposure. Single-cell mechanical testing by atomic force microscopy (AFM) showed a significant increase in the stiffness of samples following Pb^2+^ uptake (Fig. 5B). However, when yeast biomass is treated with solutions containing higher Pb^2+^ levels, mechanical stiffness is not noticeably increased (*C*_*0*_ 500 *vs* 1,000 ppb Pb^2+^). This cannot be attributed to a potential saturation of the yeast cell wall binding sites, as Pb^2+^uptake increases significantly with the increase of *C*_*0*_ from 500 to 1,000 ppb Pb^2+^ (Fig. 2E). While the exact mechanism for this stiffness change is yet to be determined, it is possible that adsorption of even a thin layer of Pb^2+^, as evidenced by the TEM data (Fig. 3D), can act as a film on the cell surface that fuses the fibrillar structures together, which then effectively resists deformation more than the untreated cell wall.

## Discussion

This work reports the biosorption isotherm and kinetics of initial Pb^2+^ concentration at the ppb scale, using lyophilized *S. cerevisiae* yeast cells as biosorbents. Comparing our results with prior studies of similar systems at the ppm scale it can be concluded that the biosorption processes at the ppb scale happen faster; the fastest equilibrium attainment reported at the ppm scale is 10 min(*18*), while our results show that equilibrium is achieved in the first 5 minutes of contact. The adsorption isotherm reported in our study (Fig. 2E) follows the same pattern as the adsorption isotherms reported in the ppm literature (*e*.*g*., fig. S1). Interestingly, the maximum Pb^2+^ uptake capacity of 12 mg/g reported in this study is in the same range as the uptake capacities reported in the ppm literature for untreated inactive *S. cerevisiae* yeast, *i*.*e*., 2-30 mg/g (*e*.*g*., table S3), proving the suitability of this biomaterial as a biosorbent at the ppb scale. pH is a significant factor in the biosorption process in both scales. The rapid biosorption and high Pb^2+^ uptake are advantageous for the large-scale application of this inexpensive and abundant biomaterial suitable for the removal of trace heavy metals from water.

From the performed analyses, it can be concluded that the cell wall of *S. cerevisiae* contributes significantly to Pb^2+^ biosorption, and in particular its carboxylic acid and amide groups. By excluding chitin as a biosorbent, mannoproteins and β-glucans are the potential key *S. cerevisiae* cell wall components, which should be further analyzed to elucidate the biosorption mechanisms involved. The combined outcomes of the TEM, the spectroscopic analyses, and the cellular nanomechanical characterization, validate the likelihood of N-linked σ-hole attraction to Pb^2+^ species as a possible mechanism of biosorptive Pb retention by the mannoprotein/β-glucan cell wall fraction(*33, 34*), leading to supramolecular assemblies that make yeast cells stiffer after biosorption. These findings open new experimental pathways for approaching the challenging task of biosorption investigation at the ppb scale.

The results showed herein, together with the fact that 3 million tons of yeast are used annually by the global fermentation industry(*35*) and that the yeast market is expected to grow by 35% in the next 5 years(*36*), indicate that exploiting this biosorbent is practically feasible and economically attractive. In addition, due to its simplicity, this approach can be easily reproduced, locally sourced, and applied at scale. The approach described here compares favorably to many of the highly sophisticated synthetic biology and advanced nanomaterials approaches that have also been examined as candidates for heavy metal removal from water(*37, 38*). Applying such a low-value resource to remove trace contaminants from water could lead to significant environmental benefits for the water and wastewater treatment utilities, including their decarbonization due to limited energy use, and waste reduction as yeast cells are biodegradable. Moreover, potential desorption processes would allow for heavy metals reclamation, enhancing the application of circular economic models.

## Acknowledgments

We thank the MIT Koch Institute’s Robert A. Swanson (1969) Biotechnology Center for technical support, and specifically the Nanotechnology Materials Lab for assisting in TEM sample preparation and imaging. We acknowledge the MIT Materials Research Science and Engineering Center (MRSEC) for assisting with TEM imaging and ATR-FTIR analyses. We also thank the MIT Center for Environmental Health Sciences (CEHS) and in particular Dr. Bogdan Fedeles for the helpful discussions and troubleshooting concerning the ICP-MS analyses of our samples. We are grateful to Dr. Constantinos Katsimpouras from the MIT Metabolic Engineering Laboratory for assisting with the HPLC analyses. We also thank Lorena Altamirano for her assistance with biological methods and protocols.

## Author contributions

Conceptualization: P.M.S., C.E.A., M.T., H.G., N.G.

Methodology: P.M.S., C.E.A., M.T., J.G., E.M.D.

Investigation: P.M.S., C.E.A. J.G., C.B.

Visualization: P.M.S., C.E.A.

Funding acquisition: P.M.S., C.E.A., F.T., A.M., N.P.P., B.W.S., N.G.

Project administration: P.M.S., C.E.A.

Supervision: C.E.A., M.T., H.G., N.G.

Writing – original draft: P.M.S., C.E.A.

Writing – review & editing: All authors

## Data and materials availability

All data are available in the main text or the supplementary materials. Additional data supporting the results presented herein can be provided upon request.

