## Supplementary Materials for "Investigating lead removal at trace concentrations from water by inactive yeast cells"

Materials and Methods

Glassware cleaning

Glassware was autoclaved for 15 minutes at 121°C (tabletop autoclave 3870E, Tuttnauer, USA) and rinsed three times with ultrapure, type 1 ultrapure water with a resistivity of 18.2 MΩ.cm at 25 °C and TOC<5 ppb (Milli-Q Direct 8 Water Purification System, MilliporeSigma, USA). Phosphate-free detergent suitable for trace heavy metals analyses was used to wash the glassware (Liquinox detergent, Alconox, USA), which was then rinsed 3 times with Milli-Q water and soaked in a 20% nitric acid (HNO_3_) bath for at least 24 hours. The nitric acid bath was made using 69% HNO3 (ARISTAR PLUS, VWR Chemicals BDH, VWR, USA) and Milli-Q water. After soaking in the 20% HNO_3_ bath, flasks were rinsed 3 times with Milli-Q water prior to use.

Yeast strain, culture & OD600 culture density measurements

The *S. cerevisiae* Meyen ex E.C. Hansen MYA-796 strain was used (ATCC, Virginia, USA). YM agar and broth media (ATCC 200 YM Medium, ATCC, Virginia, USA) were used for the yeast cultures. Yeast cells were incubated (Multitron Standard incubator shaker, INFORS HT, USA) at 30 ^o^C and 200 rpm in 2 L Erlenmeyer flasks with shallow medium content (100-200 mL). Discrete time-point optical density measurements (OD) were performed using 2 mL non-frosted cuvettes (VWR Standard Spectrophotometer Cuvettes, VWR, USA) and a tabletop, ultraviolet-visible spectrophotometer measuring at 600 nm (NanoPhotometer NP80, Implen, Munich, Germany).

Biomass harvesting, washing & lyophilization

Yeast cells were harvested by centrifugation at 1 g/2,000 rpm for 10 mins (Allegra X15-R Centrifuge, USA) and washed by two successive suspensions and centrifugations with ultrapure Milli-Q water. To decide the number of washes required to remove medium residues and metabolites from cells, supernatant samples after each wash were analyzed using high-performance liquid chromatography (Aminex HPX-87H Column, Bio-Rad, USA). Harvested, washed cells were kept at -30 ^o^C for 24 hours and then inserted in the freeze dryer (Freezone 6 Liter Manifold, Labconco, USA). Each lyophilization cycle (temperature < -40 ^o^C, pressure < 0.371 mbar) lasted for at least 50 hours. Lyophilized cells (powder form) were stored in a desiccator containing silica gel.

Aqueous solutions preparation

Type 1 Milli-Q water spiked with lead(II) nitrate (Pb(NO_3_)_2_) (Sigma-Aldrich, MiliporeSigma, USA) to achieve initial Pb concentrations of up to 1,000 ppb. The initial solution pH was adjusted to the required values by using 69% HNO_3_ (ARISTAR PLUS, VWR Chemicals BDH, VWR, USA) and 0.1 M sodium hydroxide (NaOH) (Carolina Biological, USA). The pH of aqueous solutions was measured using a glass-body electrode suitable for ion-weak samples (InLab Pure Pro-ISM, Mettler Toledo, USA).

Biosorption experiments

All biosorption experiments were conducted in batch contact environments using 2 L Erlenmeyer flasks with lyophilized yeast cells added to 200 mL of Pb^2+^ containing aqueous solutions. Flasks were incubated at 200 rpm and 25 ^o^C. After the required contact time, yeast biomass was separated from the aqueous solution by centrifugation at 1 g/2,000 rpm for 10 mins (Allegra X15-R Centrifuge, USA). The supernatants were analyzed using inductively coupled plasma mass spectroscopy (ICP-MS) (7900 ICP-MS system, Agilent, USA) to measure the residual Pb^2+^ concentrations following standard operating procedures (Pb calibration standards and Bismuth internal standard, Agilent, USA). All experiments were carried out in triplicates and mean values are reported.

For all experiments, a control sample of just Milli-Q water spiked with Pb(NO_3_)_2_ was measured to act as a reference for the initial Pb^2+^ concentration, *C_0_*, in the solution. Pb^2+^ removal measurements were calculated by taking the ICP-MS measurements of the supernatants and subtracting from this reference to give the quantity of metal adsorbed on yeast biomass. Type 1 Milli-Q water alone was also tested via ICP-MS to make sure that there was no Pb^2+^ present in the aqueous matrix. The amount of Pb^2+^ uptake from the yeast biomass was calculated using Equation 1:

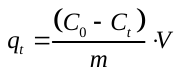
(1)

Where: *q_t_* is the Pb^2+^ mass (μg) adsorbed per gram of yeast biomass after t contact time (μg/g); *C_0_* is the initial Pb^2+^ concentration in the aqueous solution (μg/L or ppb); *C_t_* is the residual Pb^2+^ concentration measured after t contact time (μg/L or ppb); *m* is the dry weight of yeast biomass in the solution (g), and *V* is the volume of the aqueous solution (L). If equilibrium has been reached and *C_t_* is *C_e_*, then *q_t_* is *q_e_* (*17*).

Using the experimental Pb^2+^ biosorption isotherm data, *i.e.,* the *q* values at equilibrium measured after the 1 h contact of yeast biomass (*m*: 0.005 g) with aqueous solutions (0.2 L) of different initial Pb^2+^ concentrations (*C_0_*: 20, 40, 100, 200, 300, 500, 700 and 1,000 ppb), the adsorption equilibrium isothermal model was developed. The data are accurately fitting with the Langmuir adsorption isotherm model(*23*) [R^2^: 0.98], expressed by Equation 2:

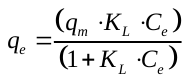
(2)

Where: *q_e_* is the Pb^2+^ uptake by yeast biomass at equilibrium (μg/g); *q_m_* is the maximum adsorption capacity estimated by the Langmuir model (mg/g); *K_L_* is the ratio of the adsorption rate and desorption rate (L/mg); and *C_e_* is the residual Pb^2+^ concentration at equilibrium (μg/L or ppb)(*24*).

Biosorption experiments and analyses using chitin were conducted following the same steps, procedures, and instruments described above, using chitin instead of yeast biomass (Chitin from shrimp shells, Sigma-Aldrich, MiliporeSigma, USA).

SEM sample preparation and analysis

After biosorption, yeast biomass was harvested via centrifugation at 1 g/2,000 rpm for 10 mins (Allegra X15-R Centrifuge, USA) and lyophilized, following the procedures described above. Samples of lyophilized yeast biomass (before and after Pb^2+^ biosorption) were coated in a 20 nm layer of gold (150TES Sputter Coater, EMS, USA), mounted onto carbon double-sided tape, and imaged with an SEM using secondary electron detection at 10 kV beam voltage and varying magnifications (FlexSEM 1000, Hitachi, Tokyo, Japan).

TEM sample preparation and analysis

Harvested yeast cells were fixed with 2% glutaraldehyde and 2.5% paraformaldehyde in 100 mM sodium cacodylate buffer (EMS, USA) for 1 h at 4 ^o^C. Then cells were washed twice with 100 mM sodium cacodylate buffer, each time re-suspending them by vortexing, storing them at 4 ^o^C for 15 mins, and centrifuging them for 10 mins at 1,000 g/2,070 rpm. Then cells were resuspended in 100 mM sodium cacodylate buffer and stayed overnight at 4 ^o^C. Pelleted cells were post-fixed for 30 min at 4 ^o^C with 1% osmium tetroxide in 0.1 M imidazole of pH 7.5 (EMS, USA). Osmium post-fixation was followed by washing the cells three times with 100 mM sodium cacodylate buffer, storing them each time for 15 mins at room temperature. The cells were then washed three times with 0.05 M maleate buffer of pH 5.15 (EMS, USA), storing them each time for 15 mins at room temperature. The cells were then stained in 2% uranyl acetate (EMS, USA) overnight, in the dark at room temperature. Stained cells were washed three times with 0.05 M maleate buffer of pH 5.15, storing them each time for 15 mins at room temperature. Then, the cells were serially dehydrated using 30%, 35%, 50%, 70%, 90%, 95%, and 100% ethanol solutions. Dehydrated cells were embedded in EMBED-812 resin (EMS, USA). Sections were cut using an ultramicrotome (EM UC7, Leica, Wetzlar, Germany) and a Diatome diamond knife at a thickness setting of 50 nm. The sections were examined using a TEM (Tecnai G2 Spirit TWIN, FEI, USA) at 120 kV.

Measurement of yeast cell wall dimensions

Yeast cell walls were measured using Image J software, taking the average measurements of 15 control and 15 Pb^2+^-exposed cells.

ATR-FTIR analysis

Samples of lyophilized yeast biomass (before and after Pb^2+^ biosorption) were put on the ATR sampling accessory of an FTIR spectrometer (Nicolet iS50 FTIR Spectrometer, ThermoFisher Scientific, USA). The measured wavenumber range was 400−4,000 cm^-1^ and for all spectra 32 scans were recorded and averaged with a resolution of 4 cm^-1^ for each spectrum. All the handling material and the ATR sampling surface were cleaned with analytical grade isopropanol and rubbed dry with clean paper before contact with the host material.

XPS analysis

Lyophilized yeast cells were taken with a spatula and loaded on a stainless steel XPS stage (5 mm width). All samples were analyzed in the same XPS run (spot size 400 μm) (K-Alpha XPS system, ThermoFisher Scientific, USA). Survey scans used a pass energy of 50 eV, while the core scans used a pass energy of 1.5 keV and were energy-calibrated using the C-C bond energy at 284.6 eV. The three peaks observed in the C 1*s* spectrum were fitted to identify changes in the C bonds due to Pb^2+^ biosorption. Data processing and peak fitting was done using the Thermo Avantage software and the Powel fitting algorithm. All the handling material and the polyacetal surface were cleaned with analytical grade isopropanol and rubbed dry with clean paper before contact with the host material.

AFM sample preparation and analysis

Lyophilized yeast biomass (following incubation with or without Pb^2+^ at 200 rpm and 25 ºC for 1 h) was harvested via centrifugation at 1 g for 5 mins and adhered to plasma cleaned and CellTak (Corning Inc, USA) adhesive-treated plastic 50 x 9 mm petri dishes, as described elsewhere(*39*). AFM force analysis was conducted with an MFP-3D AFM (Asylum Research, USA) using spherical colloidal tip CP-PNP-Au-A probes (NanoAndMore, USA) with a sphere diameter of 2 μm and a nominal spring constant of 0.08 N/m. Cellular spring constants (k_cell_) were calculated as described elsewhere using a two-spring model(*39,40*).

Figure creation

Raw data were collected and stored in CSV or Excel file formats. Plots were using the Origin software.

Supplementary Text

Extended description of the ATR-FTIR results

Control (*C_0_*: 0 ppb Pb^2+^) and Pb^2+^-exposed freeze-dried yeast cells (*C_0_*: 100 and 1,000 ppb Pb^2+^) were analyzed *via* ATR-FTIR. In all three samples, a very strong and broad hydroxyl peak was present near 3,300 cm^-1^, representing the symmetric stretching vibration of the O-H bonds, while a sharp peak close to 1650 cm^-1^ is attributed to carbonyl groups of proteins (C=O). The presence of both the hydroxyl (O-H) and the carbonyl (C=O) peaks indicates that the yeast biomass contains the carboxylic acid functionality (COOH), participating in hydrogen bonding. Absorption bands at 2,925 cm^-1^ may result from the stretching vibrations of C–H (alkanes), as they are indicative of sp3 hybridized carbons bonded to hydrogens (Csp^3^-H).

Peaks at 2,247 cm^-1^ and 2,173 cm^-1^, which are present at the control yeast biomass, undergo a significant signal reduction after Pb^2+^ biosorption, while new peaks appear after biosorption at 2,215 cm^-1^, 2,139 cm^-1^ and 2,088 cm^-1^. In addition, the peak at 2,043 cm^-1^ becomes stronger after Pb^2+^ biosorption, while the peak at 2,036 cm^-1^ becomes weaker after biosorption. All these peaks represent C≡N and C≡C stretches, indicating that these bonds contribute to the biosorption process. A blue peak shift was observed after biosorption from 1,538 cm^-1^ (control) to

1,530 cm^-1^. This shift corresponds to N-H in-plane bending from secondary protein amides, which overlaps with the C–N and NO_2_ asymmetric stretching. In addition, a blue peak from 1,397 cm^-1^ (control) to 1,383 cm^-1^ was noticed, corresponding to vibrational changes of the C-N amide group, or of the N-O functional group. Furthermore, a blue peak shift from 1,241 cm^-1^ to 1,234 cm^-1^ corresponds to C–O stretching in the esters and carboxylic acid groups. These changes, which are collectively presented in Table S1, indicate the contribution of amide and carboxylic acid groups to Pb^2+^ biosorption and the potential role of N in the yeast cell wall on Pb^2+^ binding(*41,42*).

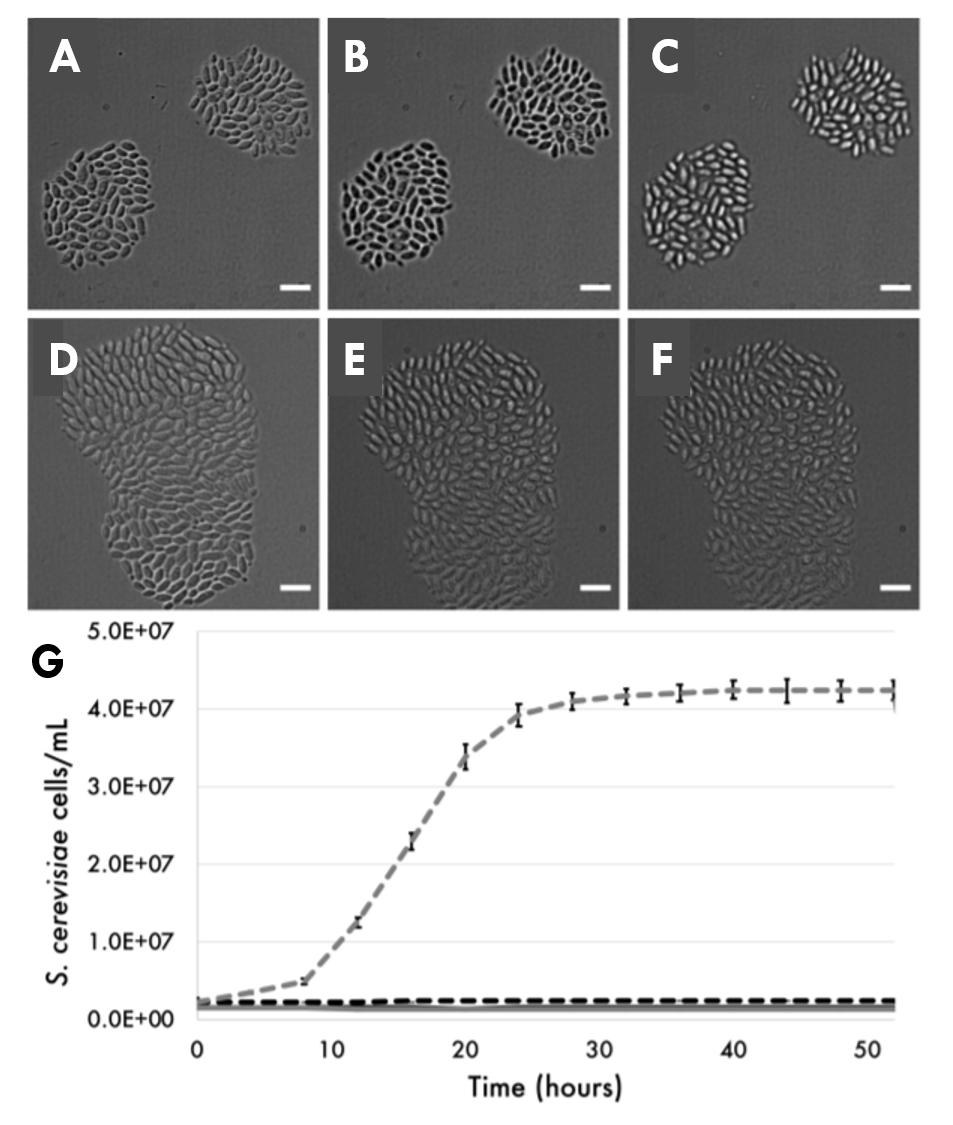

Fig. S1.

**Growth analysis of lyophilized yeast cells.** (**A**) Time course micrographs of lyophilized *S. cerevisiae* incubated in water at 0 h (scale bar: 10μm). (B) Time course micrographs of lyophilized *S. cerevisiae* incubated in water at 24 h (scale bar: 10μm). (C) Time course micrographs of lyophilized *S. cerevisiae* incubated in water at 48 h (scale bar: 10μm). (D). Time course micrographs of lyophilized *S. cerevisiae* incubated in YM growth medium at 0 h (scale bar: 10μm). (E) Time course micrographs of lyophilized *S. cerevisiae* incubated in YM growth medium at 24 h (scale bar: 10μm). (F) Time course micrographs of lyophilized *S. cerevisiae* incubated in YM growth medium at 48 h (scale bar: 10μm). (G) Growth curve obtained from OD600 measurements of lyophilized *S. cerevisiae* incubated in water (solid black line) and YM growth medium (solid gray line) and of non-lyophilized *S. cerevisiae* in water (dashed black) and YM growth medium (dashed gray) incubated at 25ºC and 200 rpm; error bars: ± 1 s.d.

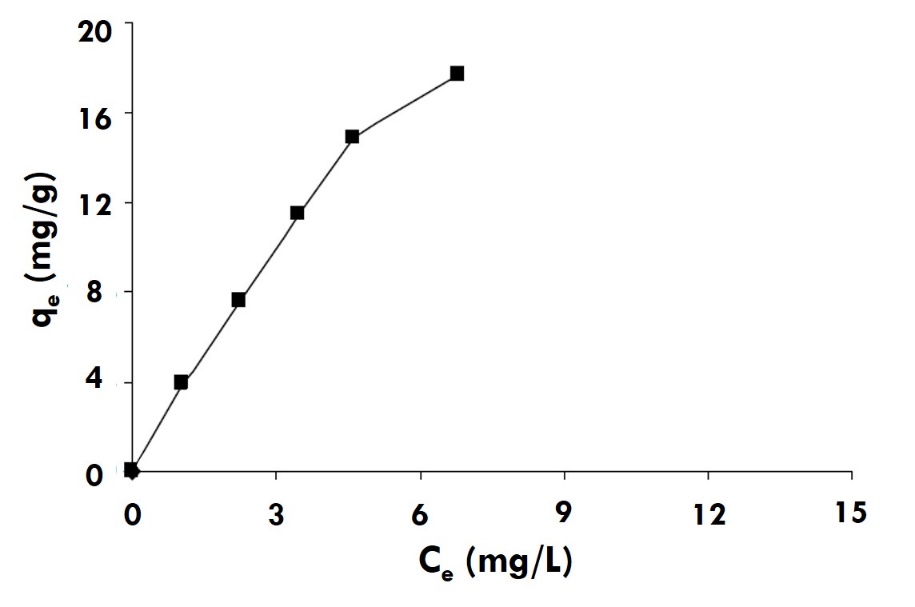

Fig. S2.

**Adsorption isotherm of Pb^2+^ by ethanol treated baker’s yeast in the ppm scale (*26*)**.

Table S1.

Changes in peaks before and after Pb^2+^ biosorption, as observed by the ATR-FTIR analysis; Control: Cells before biosorption; 100 ppb: Cells after biosorption in aqueous solutions with *C_0_* 100 ppb Pb^2+^; 1,000 ppb: Cells after biosorption in aqueous solutions with *C_0_* 1,000 ppb Pb^2+^.

| **Wavenumber** | | | **Vibration / functional groups** | **Comments** |
| --- | --- | --- | --- | --- |
| **Control** | **100 ppb** | **1,000 ppb** |  |  |
| 2,247 cm^-1^ | - | - | C≡N stretch | Peak disappeared after biosorption |
| - | 2,215 cm^-1^ | 2,215 cm^-1^ | C≡C stretch | New peak appeared after biosorption |
| 2,173 cm^-1^ | - | - | C≡N, C≡C stretch | Peak disappeared after biosorption |
| - | 2,139 cm^-1^ | 2,139 cm^-1^ | C≡C stretch | New peak appeared after biosorption |
| - | 2,088 cm^-1^ | 2,088 cm^-1^ | C≡N, C≡C stretch | New peak appeared after biosorption |
| 2,043 cm^-1^ | 2,043 cm^-1^ | 2,043 cm^-1^ | C≡N, C≡C stretch | Peak became stronger after biosorption |
| 2,036 cm^-1^ | 2,036 cm^-1^ | 2,036 cm^-1^ | C≡N, C≡C stretch | Peak became weaker after biosorption |
| 1,962 cm^-1^ | - | - | C–H overtone, weak (aromatic compound) | Peak disappeared after biosorption |
| 1,741 cm^-1^ | - | - | C=O stretch (aliphatic) from esters group | Peak disappeared after biosorption |
| 1,538 cm^-1^ | 1,531 cm^-1^ | 1,530 cm^-1^ | N-H in plane bend; NO_2_ asymmetric stretch | Peak shifted after biosorption |
| 1,397 cm^-1^ | 1,383 cm^-1^ | 1,384 cm^-1^ | C-H bend; C-N stretch; NO_2_ symmetric stretch;  N-O stretch | Peak shifted after biosorption |
| 1,241 cm^-1^ | 1,234 cm^-1^ | 1,234 cm^-1^ | C-N stretch (aliphatic amines) | Peak shifted after biosorption |
| 854 cm^-1^ | - | - | C=C bending | Peak disappeared after biosorption |
| 532 cm^-1^ | - | - | PO_4_^-3^ bend | Peak disappeared after biosorption |

Table S2.

Surface atomic concentrations of yeast cells before and after Pb2+ biosorption, as measured during the XPS analysis.

| **Element** | **Atomic concentration (%)** | | |
| --- | --- | --- | --- |
|  | **Control cells** | **Cells after biosorption**  **(*C_0_*: 100 ppb Pb^2+^)** | **Cells after biosorption**  **(*C_0_*: 1,000 ppb Pb^2+^)** |
| Carbon | 70.9% | 71.6% | 62.2% |
| Oxygen | 27.8% | 26.9% | 36.2% |
| Nitrogen | 1.3% | 1.5% | 1.6% |

Table S3.

XPS peaks profile data for yeast cells before and after Pb^2+^ biosorption.

| **Peak** | **Binding energy (eV)** | **Full width at half maximum - FWHM (eV)** | **Area (P) CPS.eV** |
| --- | --- | --- | --- |
|  | ***Control cells*** | | |
| C-C | 284.3 | 1.5 | 142,838 |
| C-N | 285.9 | 1.3 | 98,326 |
| O-C=O | 287.5 | 1.5 | 36,566 |
| C=O | 288.6 | 0.9 | 3,442 |
|  | ***Cells after biosorption (C_0_: 100 ppb Pb^2+^)*** | | |
| C-C | 284.3 | 1.9 | 72,014 |
| C-N | 285.9 | 1.1 | 67,571 |
| O-C=O | 287.5 | 2.1 | 68,559 |
| C=O | 288.6 | 1.5 | 3,204 |
|  | ***Cells after biosorption (C_0_: 1,000 ppb Pb^2+^)*** | | |
| C-C | 284.3 | 2.1 | 17,129 |
| C-N | 285.9 | 1.2 | 7,227 |
| O-C=O | 287.5 | 2.2 | 9,679 |
| C=O | 288.6 | 1.4 | 627 |

Table S4.

Prior studies on Pb biosorption by inactive yeast at the ppm scale.

| **#** | **Key functional groups involved** | **C_0_ [ppm]** | **pH** | **Incubation conditions** | **Time needed to attain equilibrium** | **Biomass used** | **Max measured q (mg/g)** | **Source** |
| --- | --- | --- | --- | --- | --- | --- | --- | --- |
| 1 | - | 100 | - | 30 °C; 140 rpm | 10 min | Commercial dry baker’s yeast washed 5 times with pure water, fixed with  ethanol & phosphorylated | ~190 | (*18*) |
| 2 | - | 200 | 5 | 25 ^o^C; 150 rpm | 15 min | EDTA-treated *S. cerevisiae* cells, oven-dried at 45°C for 24 h | 99 mg/g | (*43*) |
| 3 | C=O, OH,  NH, protein amide II band, PO_2_-, mannans, sulfur &  sulfur-oxygen compounds | 300 | 5 | 28 °C; 150 rpm | 20 min | *S. cerevisiae* - Assuit University Mycological Centre 3875, washed twice with distilled water & autoclaved | 30 mg/g | (*44*) |
| 4 | COOH, C=O, C-O & NH | ~25 | 5 | 25 ^o^C; 150 rpm | - | *S. cerevisiae* from brewing industry, washed, oven-dried at 60^o^C and ground | - | (*45*) |
| 5 | carboxylic acids & amines | 100 | 5 | 30^o^C; 150 rpm | > 60 min | *S. cerevisiae* from a brewery in India; cells washed 3 times with double distilled water, oven-dried at 80^o^C for 8 h | 2.34 mg/g | (*46*) |
| 6 | - | 207 | - | - | 2 h | *S. cerevisiae* washed | - | (*47*) |
| 7 | - | 25 | 5 | 30 ^o^C; 200 rpm | >15 min | Waste *S. cerevisiae* obtained from yeast company at Izmir; Caustic- & ethanol-treated, dried at 70^o^C for 12 h and then ground | 17.5 mg/g | (*26*) |

Movie S1.

Time lapse phase-contrast microscopy video showing the growth of lyophilized *S. cerevisiae* cells in water - no cell growth or division observed during the 52-h time course (scale bar: 10μm).

Movie S2.

Time lapse phase-contrast microscopy video showing the growth of lyophilized *S. cerevisiae* cells in growth media - no cell growth or division observed during the 52-h time course (scale bar: 10μm).
